# Competition between tool and hand motion impairs movement planning in limb apraxia

**DOI:** 10.1101/2025.04.07.647589

**Authors:** Simon Thibault, John B. Yates, Laurel J. Buxbaum, Aaron L. Wong

## Abstract

Tool use is a complex motor planning problem. Prior research suggests that planning to use tools involves resolving competition between different tool-related action representations. We therefore reasoned that competition may also be exacerbated with tools for which the motions of the tool and the hand are incongruent (e.g., pinching the fingers to open a clothespin). If this hypothesis is correct, we should observe marked deficits in planning the use of incongruent as compared to congruent tools in individuals with limb apraxia following left-hemisphere stroke (LCVA), a disorder associated with abnormal action competition. We asked 34 individuals with chronic LCVA (14 females) and 16 matched neurotypical controls (8 females) to use novel tools in which the correspondence between the motions of the hand and tool-tip were either congruent or incongruent. Individuals with LCVA also completed background assessments to quantify apraxia severity. We observed increased planning time for incongruent as compared to congruent tools as a function of apraxia severity. Further analysis revealed that this impairment predominantly occurred early in the task when the tools were first introduced. Lesion-symptom mapping analyses revealed that lesions to posterior temporal and inferior parietal areas were associated with impaired planning for incongruent tools. A second experiment on the same individuals with LCVA revealed that the ability to gesture the use of conventional tools was impaired for tools rated as more incongruent by a normative sample. These findings suggest that tool-hand incongruence evokes action competition and influences the tool-use difficulties experienced by people with apraxia.

**Significance Statement:** Prior research indicates that competition between different representations associated with moving or using tools must be resolved to enable tool use. We demonstrated that competition may be exacerbated when tool and hand motions are incongruent (e.g., pinching the hand opens a clothespin), resulting in tool-use impairments particularly for individuals with greater severity of limb apraxia, a disorder known to be associated with action competition abnormalities. Lesions in posterior portions of the brain’s tool use network were associated with impairments in planning incongruent tool actions. This study thus demonstrates that tool-hand incongruence may invoke competition between motions of the hand and tool-tip, which individuals with limb apraxia have difficulty resolving to properly use tools.

## Introduction

Tool use enables interactions with our environment and is associated with complex motor planning demands. While many tools can be treated as simple extensions of the hand (e.g., the toothbrush and hand simultaneously move horizontally or vertically), other tools may pose a greater challenge for movement planning because they are associated with incongruent motions of the tool and the hand (i.e., clothespins open by pinching the fingers closed). In neurotypical individuals, incongruent tools have been associated with longer planning times compared to congruent tools (Massen and Prinz, 2007; Beisert et al., 2010; Baugh et al., 2012). This increased planning difficulty may arise from a need to resolve competition between the different motions of the tool tip and the hand. Such competition could emerge because people can represent hand and tool motions in an abstract fashion (e.g., desired motion paths through space; Wong et al., 2016, 2019). The motion path of the tool is automatically activated when preparing to use it (e.g., opening motion; Beisert et al., 2010; Zhang et al., 2021). When this motion is incongruent with the required hand motion, a second motion representation also becomes activated (e.g., pinching). The result is two motion representations that compete for selection: an automatic congruent motion (opening) and a goal-directed, incongruent one (pinching; Hardwick et al., 2019; Zhang et al., 2021). Resolving this competition to select the incongruent motion representation for controlling the hand may require additional response time and opportunity for errors. On the other hand, a recent study found that people were not slower to plan tools with incongruent tool-hand motions (versus congruent ones; Ras et al., 2022), suggesting that these motion representations may not actually compete. Thus, it remains an open question whether incongruent tools pose greater demands on our planning ability.

One way to address this question is to examine individuals with limb apraxia following a left hemisphere stroke (LCVA), which can markedly affect tool-related actions (Buxbaum and Kalénine, 2020). Prior studies in this population have identified a form of competition between different action representations associated with moving or using tools (e.g., power grasp versus non-prehensile “poke” for a calculator; Watson and Buxbaum, 2015; Garcea and Buxbaum, 2023). Impaired competition resolution slows planning times and reduces gesture accuracy in apraxic individuals (Jax and Buxbaum, 2013; Lee et al., 2014; Bolognini et al., 2015; Garcea et al., 2019). Thus, studying how people with apraxia plan tool-use actions serves as a sensitive assay of underlying action competition. However, it remains unclear if these action-competition deficits also impact the ability of apraxic individuals to plan actions with incongruent tools. Impairments in resolving tool-use competition have been associated with lesions to the left supramarginal gyrus (SMG), inferior frontal gyrus (IFG), and insula (Watson and Buxbaum, 2015; Garcea et al., 2019; Garcea and Buxbaum, 2023). If incongruent tools also invoke action competition, similar brain regions may support such tools. Neuroimaging studies suggest that the use of incongruent tools may indeed activate the SMG, but also the posterior temporal cortex (pTC; Gallivan et al., 2013; Ras et al., 2022). Thus, the brain regions supporting tool-related competition resolution remain unclear Here we examined the relationship between limb apraxia and the ability to plan the use of both novel and conventional incongruent tools. In the first experiment, we asked participants with LCVA with varying degrees of apraxia to use congruent and incongruent novel tools. These unfamiliar tools allowed us to isolate the effect of tool-hand incongruence on planning independent of any prior tool-use knowledge. In the second experiment, we examined how the ability of these same participants to gesture the use of conventional tools varied as a function of tool-hand incongruence. We predicted in both experiments that if incongruent tool-hand motion invokes action competition, tool-use impairments would be affected by the degree of tool-hand incongruence and the severity of apraxia. We also performed exploratory lesion-symptom mapping analyses to identify brain regions critical for resolving tool-hand incongruence.

## Materials and Methods

### Experiment 1: Effect of tool-hand incongruence on novel tool-use planning

#### Participants

Fifty-six participants completed this study, including 37 individuals with chronic left hemisphere cerebral vascular accident (LCVA) and 20 neurotypical controls. Four neurotypical controls were excluded for failure to achieve a score above the normative cutoff for mild cognitive impairment (≥24; see Carson et al., 2018; Milani et al., 2018) on the Montreal Cognitive Assessment (MoCA) or to understand and follow task instructions. Two people with LCVA were excluded because brain scans collected after behavioral testing revealed that their stroke was not strictly confined to the left hemisphere. A third person with LCVA was excluded because they were identified as being left-handed prior to their stroke according to the Edinburgh Handedness Inventory (Oldfield, 1971). The final sample included 50 right-handed participants (34 people with LCVA and 16 neurotypical controls). To mitigate any possible impact of hemiparesis in the right arm, all participants (including neurotypicals) performed the task with their non-dominant left hand. The two groups did not significantly differ in age, education and gender (Table 1). All participants were recruited through the Research Registry of the Jefferson Moss Rehabilitation Research Institute. Study procedures were approved by the Thomas Jefferson University Institutional Review Board. Participants provided written informed consent and were compensated for their time.

**Table 1:** Demographics for the sample tested in Experiment 1. To ensure that the groups did not differ significantly in age and education, two sample t-tests were conducted. A chi-squared test was run to test whether the proportion of females and males was statistically different across groups. Demographics are reported under the form Mean ± Standard Deviation.

|  | Neurotypical Controls (N = 16) | LCVA (N = 34) | Statistics |
| --- | --- | --- | --- |
| Age (years) | 62.81 ± 10.89 | 63.65 ± 10.21 | $t_{(48)} = 0.26$; $p = 0.79$ |
| Education (years) | 15.69 ± 2.33 | 14.76 ± 2.87 | $t_{(48)} = 1.12$; $p = 0.26$ |
| Sex | 8 females<br>8 males | 14 females<br>20 males | $\chi^2_{(1)} = 0.34$; $p = 0.55$ |
| Stroke onset (months) | NA | 132.72 ± 70.22 | NA |
| MoCA score | 27.50 ± 1.26 | NA | NA |

#### Materials

We designed and built a set of six novel tools to isolate the effect of tool-hand incongruence on movement planning and avoid confounds due to potential variability in tool familiarity and tool-use knowledge. Each tool was associated with the same task objective: to push a tall cylindrical object (3D-printed and weighted to resist toppling) into a target zone. Each tool had a “body” made of wood, a designated handle, and a tool-tip comprised of a 3D-printed bracket that fit snugly around the cylinder (Fig. 1). Each tool was also associated with its own workspace, upon which was drawn a square denoting the target region. Two pairs of tools were constrained by a pivot and required a rotation movement to use them. One pair of tools was congruent (Fig. 1A), as the tip of the tool and the hand moved in the same direction, and the other pair was incongruent (Fig. 1B), as the hand and tool-tip moved in opposite directions (e.g., when the hand moved abductively, the tool-tip moved adductively). The set also included a pair of unconstrained (i.e., stick-like) tools (Fig. 1C), for which people with apraxia have previously demonstrated difficulties (Jacobs et al., 2009). These tools also added variability in the types of responses participants were required to make across trials. Each pair of tools were identical except for the design of the tool body; one tool of each pair included an extension of the tool body that created a spatial offset (displacement) between the line of action of the hand and that of the tool tip. We included displaced tools to potentially exaggerate the degree of tool-hand incongruence; however, this spatial offset was not found to be associated with disproportionate difficulty in tool-use planning as compared to non-offset tools (χ^2^s < 1.75, ps > 0.18). Thus, we collapsed our data across this parameter in the reported analyses.

**Figure 1:**
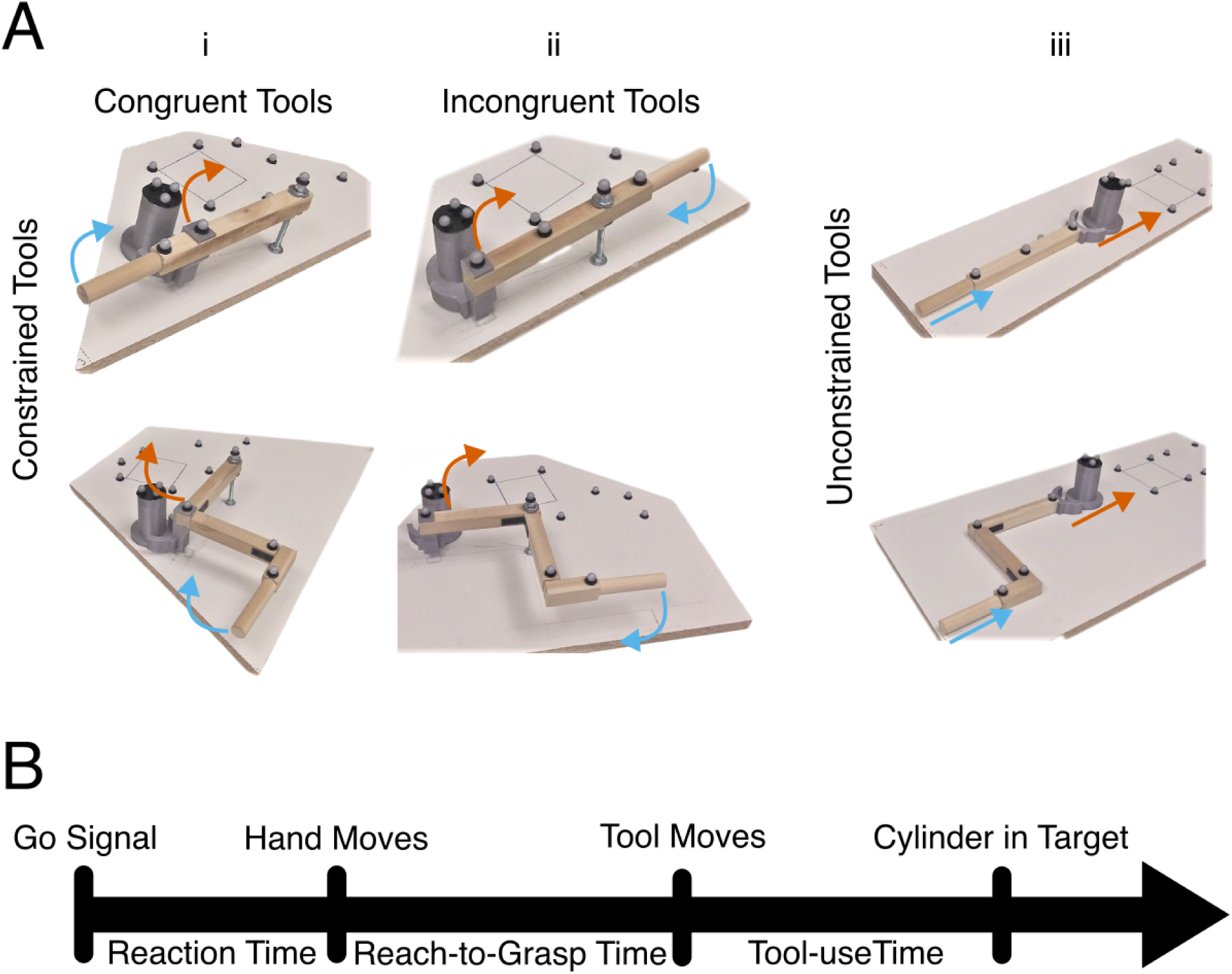
Tool set and trial timeline. (A) Set of tools used for Experiment 1: i) constrained congruent tools, for which the direction of motion of the hand and tool-tip was the same; (ii) constrained incongruent tools, for which the direction of motion of the hand and tool-tip differed; (iii) unconstrained tools lacking a pivot point. In all panels, the gray cylinder was the recipient object to be pushed into a goal, depicted as a square drawn on the table surface. The blue arrow depicts the motion of the hand acting on the tool, and the orange arrow depicts the motion of the tool tip. (B) Timeline for a single trial. At the Go signal, an opaque face mask became transparent, and participants were able to view the tool. This was followed by the initiation of hand movement, the onset of tool movement, and finally (on successful trials) the propulsion of the cylinder into the target goal. These events effectively divided the trial into 3 key time periods for analysis purposes.

Movement kinematics were recorded with a 10-camera Vicon system (Vicon, Oxford, UK) which used infrared light to track passive spherical markers. The positions of the left hand, the tools, the recipient object (cylinder), and the targets were recorded using Vicon Nexus 2.0 software (Vicon, Oxford, UK) and custom-built Python scripts. Kinematics were acquired at a sampling rate of 200Hz throughout the 8 s trial duration. Any trial where the cylinder was not inside the target after the 8 s period was treated as an error.

Vision was constrained using a custom-built polymer-dispersed liquid crystal face mask that was either opaque or transparent depending on whether an electric voltage was applied to the mask. In the opaque state, vision was completely obscured, enabling control of the timing of when participants could begin planning their response. The state of the mask was controlled using the same custom Python computer program that controlled kinematic recordings.

#### Procedure

Participants sat in front of a table with their left arm resting on an armrest, with the tip of the left ring finger on a tactile landmark. Before starting the main task, all participants completed a practice phase. The first part of the practice phase consisted of pushing the cylinder into the target with their left hand for six trials to become familiar with the overall objective of the task and the timing of the availability of vision during the trial. The rest of the practice phase consisted of six additional trials, where participants pushed the cylinder into the target with a small hand-held tool comprised of a handle attached directly to a 3D-printed tool tip (i.e., lacking any tool body). Each trial began with an auditory instruction “Prepare”, followed 1 to 3 seconds later by a go signal (i.e., a single auditory beep) and the face mask becoming transparent. Participants were instructed to leave the starting position and use the tool to move the cylinder into the target as quickly as possible after the go cue. A maximum of 8 s was allotted to perform the action; after this time the participants heard two beeps indicating that the trial was over and had an additional 2 seconds to return to the start position before the face mask again obscured their vision. During the inter-trial interval, the tool and the cylinder were reset to their initial position by the experimenter. During the practice phase, the cylinder and/or the tool were presented three times on the right side of the participant, and three times on the left side, in a randomized order. This procedure aimed to add variability in the task and prevent participants from anticipating the position of the tool handle. The entire practice block was designed to ensure that participants could understand task instructions and perform simple reach-to-gasp actions without and with a tool.

After the practice phase (∼15 minutes), participants performed the main task (∼30-45 minutes). Participants completed 6 repetitions with each of the six tools, resulting in 36 trials. As with the practice phase, across the 6 repetitions, each tool was presented with its handle on the participants’ left or right side for half of the trials in random order (i.e., 3 repetitions for each). Trials were pseudo-randomized such that participants would not complete two consecutive trials with the same tool. As a control condition, six additional repetitions with each of the congruent and incongruent tools without spatial offset presented in the reverse orientation were also interleaved throughout the task to control for the fact that the direction of hand motion required to use the congruent (i.e., away from the body) and incongruent (i.e., toward the body) tools was different. The distance between the tool handle and the hand starting position was kept constant at 100 mm, regardless of the tool’s location or orientation, enabling us to compare reach-to grasp-time across tools. Participants remained naive to the six tools of the main experiment until starting the main task. During the inter-trial interval, participants’ vision was obscured while the tool was changed by the experimenter. A short break was provided halfway through the main task.

#### Assessment of limb apraxia and stroke severity

Individuals with LCVA took part in a battery of tests to evaluate their general stroke and apraxia severity. Language comprehension was assessed with the comprehension subset of the Western Aphasia Battery (WAB). People receiving a score below 4 out of 10 (moderate impairment) were excluded. Participants also completed the National Institutes of Health Stroke Scale (NIHSS) assessing overall stroke severity. In addition, the integrity of the praxis system was tested by assessing accuracy in two tests of limb apraxia. The Gesture-to-Sight task required gesturing the use of a tool in response to a picture of that tool. The Meaningless Imitation task required imitating (in a mirror-like fashion) an arbitrary gesture that was demonstrated by an actor in a video clip. In both tasks, participants were instructed to perform gestures with their left, less-affected arm. Gestures were videotaped and later coded by two raters who reached inter-rater reliability (Cohen’s Kappa ≥ 85%) on a subset of participants.

Gestures were scored on three components: hand posture (showing how the tool would be grasped), arm posture (showing which way the arm would move through space) and amplitude/timing (showing how large in space the movement should be and how many times it is typically repeated) based on detailed scoring criteria (Buxbaum et al., 2000, 2005). These component scores can be examined separately but are more commonly averaged together to form a total task score. Previous studies have suggested that the Gesture-to-Sight and Meaningless Imitation tasks rely on different underlying processes and are associated with lesions to different brain regions (Buxbaum et al., 2014; Hoeren et al., 2014). Thus, averaging together the total Gesture-to-Sight and Meaningless Imitation scores yielded a score that more completely reflects overall apraxia severity for each person with LCVA, and captures more potential types of impairments that could contribute to impaired resolution of action competition between incongruent hand and tool motions (Buxbaum et al., 2000; Dressing et al., 2021).

#### Kinematic Data Preprocessing

Using Vicon Nexus 2.0 software, we reconstructed different skeletons (i.e. sets of markers) for each tool and target, as well as the left hand and the cylinder (n = 16 skeletons). The reconstructions allowed each marker of interest to be correctly labeled with its position in millimeters (mm). In case of marker loss (e.g., if a marker was obscured from the view of the cameras), we used the interpolation tools available in Vicon. For skeletons consisting of 4 or more markers, we used the rigid body fill algorithm; for skeletons with less than 4 markers, we used the pattern fill algorithm. Finally, we used a custom-built Python script to fill in any additional missing marker positions that could not interpolated in Vicon Nexus (i.e., discrete marker losses) using linear interpolation.

From the position data, we estimated the velocities for the left hand, the tool handle and the cylinder. For the left hand and the cylinder, we calculated the centroid of the markers. For the tool handle, we considered the position of one unique marker placed at the edge of the tool handle. Position data were smoothed using a second order Savitzky-Golay filter with a window size of 31 points. The choice of window size was made after visual inspection on representative trials to confirm a reasonable correspondence between the raw and smoothed data. 3D velocity was then calculated by taking the derivative of the smoothed position data for the left hand, tool handle and the cylinder respectively using the central difference method.

As we are primarily interested in the question of how tool-hand incongruence affects the planning of tool-use actions, our analyses focused on two stages of the planning process: reaction time (RT, i.e., the time from a go-signal to initiation of a hand movement toward the tool) and reach-to-use time (i.e., the time from hand movement onset to first grasp of the tool). Reach-to-use time is of particular interest as prior work has shown that planning related to the immediate upcoming movement (in this case, the actual use of the tool) can often occur during this period instead of or in addition to the RT (e.g., Orban de Xivry et al., 2017; Kashefi et al., 2024). Movement onset for the left hand and tool handle was identified as the first time the velocity exceeded 10 mm.s^-1^, verified by visual inspection. RT was estimated as the time from the go signal to the onset of hand movement. Trials with an RT below 100 ms were excluded from further analysis (1.15% of the trials). The reach-to-use time was estimated as the time from onset of hand movement to the onset of tool-handle movement.

#### Behavioral Analyses

In R, Linear Mixed Models (LMMs) were used to analyze RT and reach-to-use time using the package *afex* (Singmann et al., 2024). To test the effect of the apraxia severity on the ability to use novel tools, we ran LMMs only on the group of people with LCVA and included apraxia severity score as a continuous regressor. Our models initially included two additional terms, NIHSS and WAB comprehension scores, as nuisance regressors to control for the potential effect of stroke severity. As these regressors did not explain a significant proportion of variance for any dependent variable tested (χ^2^s < 1.23, ps > 0.26), we excluded these terms from our models. All models (including those described below) included a random effect of participant.

Our primary hypothesis concerned the contrast between congruent and incongruent tools on planning (either RT or reach-to-use time); thus, we focused our analyses on the subset of constrained tools (for analyses regarding the effect of Constraint, see Supplementary Material). We examined a model including Incongruence and Apraxia Severity score and their interaction. If a significant effect involving Incongruence was found, we planned to run an additional model to check whether hand motion direction could potentially explain this effect. Specifically, for the trials analyzed, the congruent tools entail abductive, away-from-body movements while incongruent tools entail adductive, toward-body movements. To control for this, the study design included a small number of interleaved trials where congruent tools required abductive and incongruent tools entailed adductive movements (control condition). Therefore, when we found a significant interaction between Incongruence and Apraxia Severity, we also tested whether Apraxia Severity score interacted with the within-subject factor Hand Direction. Since we observed that the effect of incongruence was modulated by apraxia severity, we performed a post-hoc analysis to assess whether this impairment resolved with repeated exposure. We took advantage of the six repetitions of each tool to examine performance in three bins (early, middle, and late), each containing two repetitions of each tool. We examined a model including Incongruence, Apraxia Severity, and Bin and their interactions.

We also planned to test two additional factors. First, we asked if, as a group, people with LCVA planned actions with incongruent tools differently than neurotypical controls (i.e., an LMM testing for an effect of Group and/or an interaction between Group and Incongruence). Second, we examined the effect of Incongruence on the ability to use the novel tools. Use ability was quantified as the average movement velocity, calculated as the tool-handle path length divided by the tool-use time (i.e., the time from tool-handle movement onset to the time the cylinder was placed entirely within the target); this accounted for differences in path length of the tool movement. We used a LMM to test if Incongruence and Apraxia Severity (and their interaction) affected the velocity with which participants moved the tool.

For all models mentioned above, we investigated interaction effects using post-hoc tests with a Tukey correction for multiple comparisons (*emmeans* package; Lenth, 2024). If an interaction involved a continuous regressor (e.g., Apraxia Severity), post-hoc comparisons were performed using the minimum and maximum values of this regressor as reference levels (see Lenth, 2016). All the results reported for RT and reach-to-use time are expressed in milliseconds as Mean ± Standard Error of the Mean.

#### Lesion Symptom-Mapping analyses

To examine lesions associated with poor tool-use performance, we performed an exploratory Support Vector Regression-Voxel Lesion Symptom Mapping (SVR-VLSM) analysis using the SVR-VLSM Matlab toolbox (DeMarco and Turkeltaub, 2018). SVR-VLSM is a multivariate analysis technique that uses machine learning to determine the association between lesioned voxels and behavior across all voxels and participants simultaneously. Using this approach, we analyzed data from the 29 people with LCVA in our dataset for whom we were able to obtain a structural T1 MRI scan and derive a lesion map. Lesion maps were either hand-drawn or drawn with an automated algorithm (LINDA; Pustina et al., 2016). In both cases, the lesion map was checked by an experienced neurologist (H. Branch Coslett) who was blinded to the goals of the study. The lesion maps were registered into the Montreal Neurological Institute (MNI) space. Only voxels lesioned in at least 10% (i.e., 3) of participants were included in the analysis. Total lesion volume was regressed from both behavioral scores and the lesion maps. Voxelwise statistical significance was determined using a Monte Carlo style permutation analysis (10,000 iterations) in which the association between behavioral data and lesion map was randomized. SVR-VLSM analyses were run on the reach-to-use time for incongruent and congruent tools separately. We also ran a disjunction analysis to identify lesion-symptom relationships that were stronger for incongruent than congruent tools. To capitalize on the strong behavioral differences between congruent and incongruent performance observed in the first bin of the experiment, these analyses focused on data from that bin. A voxelwise threshold of P < 0.005 was applied to determine chance-level likelihood of a lesion-symptom relationship. Within this map, we applied a further cluster-size threshold to exclude clusters with less than 100 voxels. Localization of significant grey and white matter clusters was checked with the Eve Atlas (John Hopkins University; Oishi et al., 2009); https://github.com/muschellij2/Eve_Atlas).

### Experiment 2: Effect of tool-hand incongruence on conventional tool-use gesture

#### Participants

In the second experiment, we investigated whether performance impairments associated with the use of incongruent novel tools would also be observed when people with limb apraxia gestured the use of conventional tools. To test this, we used data from the Gesture-to-Sight of Tools task (see above) from the sample of 34 people with LCVA included in Experiment 1. Gesture-to-Sight is known to provide a sensitive assay of impairments in conventional tool-use behavior (Randerath et al., 2011; Baumard et al., 2014). We considered errors for each item separately for each of three components (hand posture, arm posture and amplitude/timing; Buxbaum et al., 2000, 2005).

Prior to the experiment, we estimated the degree of incongruence of each tool by collecting ratings from a large pool of neurotypical control subjects. A total of 259 English-speaking neurotypical participants took part in an online survey on Amazon Mechanical Turk. After removing participants who were determined to not be engaged in the task (see *Procedure* below), the ratings of 131 participants were retained for further analyses (age: 37.08 ± 12.40 years, education: 15.48 ± 3.53 years, sex: 71 females, 59 males, 1 undeclared). Each of the 131 participants received monetary compensation for the completion of the survey. All study procedures were approved by the Thomas Jefferson University Institutional Review Board.

#### Materials

In total, 33 tools were tested both in the Gesture-to-Sight task and the online survey. For the online survey, a picture of each tool (the same one used in the Gesture-to-Sight task) was presented at the center of the screen. Under this picture, a question about the tool was presented along with a slider bar ranging from 1 to 100. Participants used their computer mouse to move a cursor along the slider bar in response to the question. The survey was developed on Psychopy and converted into JavaScript using a Psychopy built-in function. The survey and resulting data were hosted on Pavlovia.

### Procedure

Normative participants were required to answer two questions for each tool. First, for every tool they were asked to rate their familiarity with it: “How often have you used this tool or seen someone use it?”. The slider bar was associated with two labels at the extremes, where 1 corresponded to the label “Never” and 100 to the label “All the time”. The participants were also offered the option to check a box to indicate that they did not recognize the tool. If the box was checked, this tool was not presented with the second question. The second question asked people to rate the perceived degree of incongruence associated with each tool: “How much do the hand movements to use this tool differ from the motion of the functional part of the tool?”. The extremes of the slider bar were associated with labels: 1 corresponded to the label “Not different” and 100 to the label “Completely different”. Prior to performing ratings, participants were given clear explanations of the ratings tasks and examples of congruent and incongruent tools to aid understanding.

To ensure that participants conscientiously completed the survey, 8-12 catch trials that queried the tool name (e.g., “Is this tool named a hammer?”) were inserted in pseudo-random order. The slider bar was associated with two labels at the extremes, where 1 corresponded to the label “No” and 100 corresponded to the label “Yes”. Participants were instructed to move the cursor completely to the left or right to respond. If the participants committed more than two errors on catch trials their data were excluded. This resulted in the exclusion of 115 participants. Further visual inspection of the distribution of the ratings led to the rejection of an additional 13 participants who responded to the other survey questions by placing the cursor in the same position for every presented tool. Using the data from the remaining 131 participants, average familiarity and incongruence ratings were calculated for each of the 33 tools.

#### Statistical Analyses

We conducted an item-level analysis (i.e., considering the average performance across individuals for each tool) to assess whether accuracy of the people with LCVA in the Gesture-to-Sight task for each of the three scored components (i.e., hand posture, arm posture or amplitude/timing) was sensitive to tool-hand incongruence ratings provided by the normative sample. These components revealed the errors that occurred despite people’s best attempts to plan the correct gesture (i.e., competition resolution failures). In contrast, the time between initiation of the hand and the tool movements in Experiment 1 reflects how much time people spend trying to resolve tool-hand competition and prepare their response. Thus, the dependent variables from these two experiments do not have direct correspondence. We used a binomial General Linear Mixed Model (GLMM) in R (*afex* package; Singmann et al., 2024) to assess whether the accuracy of people with LCVA to gesture the use of individual conventional tools could be explained by the degree to which that tool was rated as incongruent by the neurotypical individuals in the survey study. The model included Component as a fixed-effect factor (i.e., hand posture, arm posture, and amplitude/timing), Tool-Hand Incongruence as a continuous regressor, and a random intercept of participant. Note that this random intercept effectively accounts for differences in total performance on the Gesture-to-Sight task (e.g., a measure of apraxia severity). In contrast, here we are interested in testing for the significance of the slope denoting how accuracy when gesturing individual tools varies as a function of the average incongruence rating of those same tools, regardless of apraxia severity. Finally, we re-ran the model with the addition of survey ratings of Tool Familiarity collected from the same set of neurotypical individuals to test if familiarity with the tools could explain the effects observed with Tool-Hand Incongruence. To investigate potential interaction effects, post-hoc tests were conducted by applying a Tukey test to correct for multiple comparisons (*emmeans* package; Lenth, 2024). If an interaction involved a continuous regressor (e.g., incongruence rating), post-hoc comparisons were performed using the minimum and maximum values of this regressor as reference levels (see Lenth, 2016). Accuracy is expressed as Mean ± Standard Error of the Mean.

## Results

### Experiment 1: Effect of tool-hand incongruence on novel tool-use planning

#### Planning to use incongruent tools is impaired as a function of apraxia severity

To examine the effect of Incongruence, our primary analyses focused on our set of constrained novel tools (i.e., tools with a pivot). Participants with LCVA included individuals with a range of apraxia severity. We assessed whether the ability to plan congruent versus incongruent tools differed as a function of apraxia severity. As noted in the Methods, stroke severity or language comprehension impairment did not significantly explain the observed experimental effects in RT or reach-to-use time (χ^2^s < 1.23, ps > 0.26). For RT, incongruent tools on average were associated with longer RTs compared to congruent tools but this difference did not reach significance (Incongruent tools: 595 ± 56 ms, Congruent tools: 548 ± 63 ms; χ^2^(1) = 2.99, p = 0.08). We did not find a main effect of Apraxia Severity (χ^2^(1) = 2.00, p = 0.15), nor an interaction between Apraxia Severity and Incongruence (χ^2^(1) = 0.46, p = 0.49).

In contrast, for reach-to-use time we found a main effect of Apraxia Severity (χ^2^(1) = 10.44, p = 0.001). This effect reflects a slowing of reach-to-use time as apraxia severity increased. The main effect of Incongruence was also significant (χ^2^(1) = 4.18, p = 0.04): reach-to-use times were longer for incongruent versus congruent tools (Incongruent tools: 1031 ± 56 ms, Congruent tools: 967 ± 49 ms). Importantly, we also found a significant interaction between Apraxia Severity and Incongruence (χ^2^(1) = 3.87, p = 0.04). This interaction was driven by slower reach-to-use times for incongruent compared to congruent tools particularly for people with greater apraxia severity (post-hoc test: p = 0.01; Fig. 2A); no such difference in reach-to-use time as a function of Incongruence was observed for people with less severe apraxia (post-hoc test: p = 0.38).

**Figure 2:**
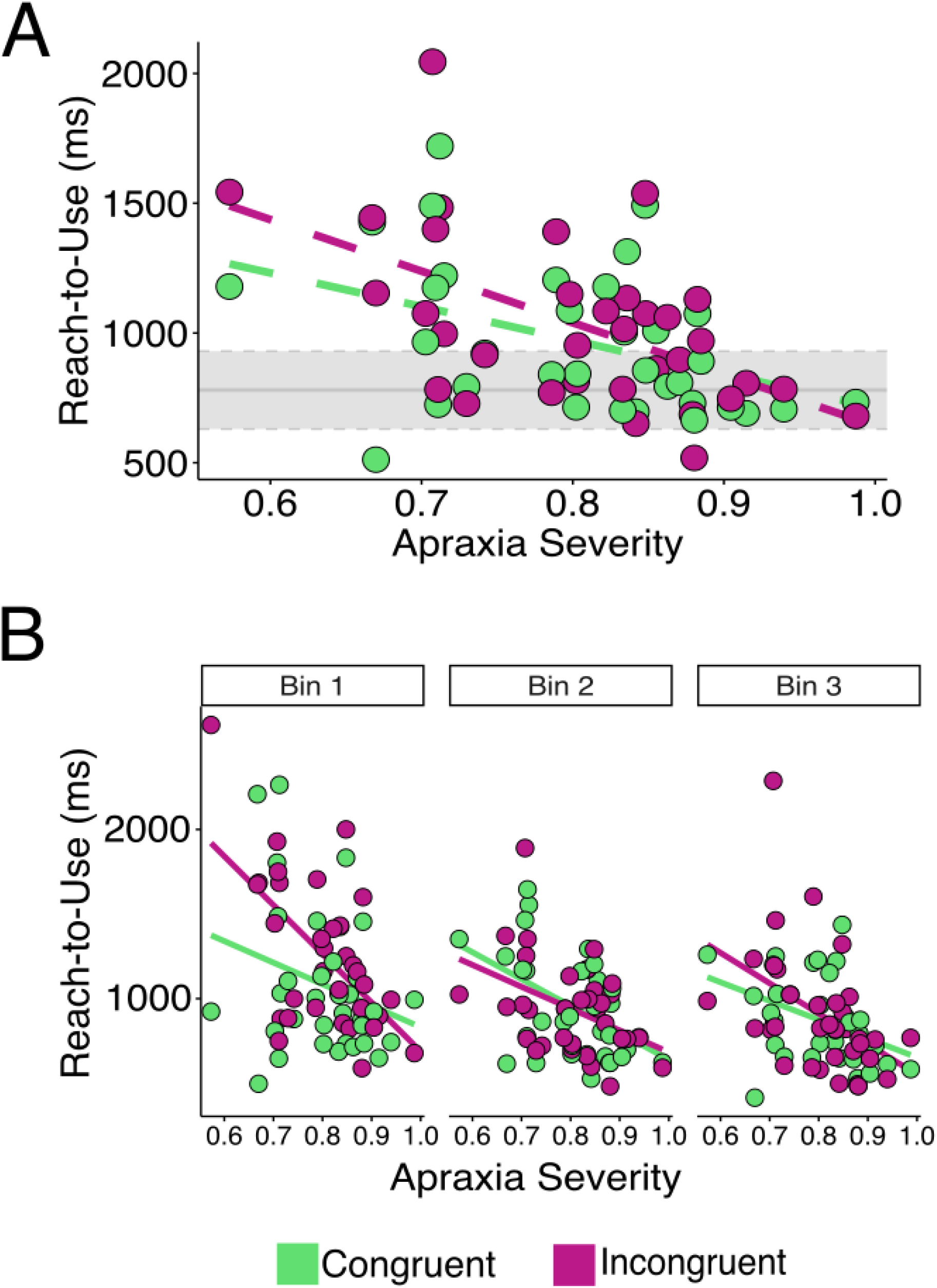
Tool-use planning deficits as a function of apraxia severity. (A) Reach-to-use time was longer for incongruent (red) as compared to congruent tools (green) as a function of apraxia severity. The horizontal axis represents apraxia severity, with lower values indicating greater severity. Each LCVA individual is represented by a unique dot for incongruent tools (red) and congruent tools (green). The neurotypicals’ performance range across both congruent and incongruent tools is depicted by the light grey area, with the solid line being the neurotypical mean performance, and the dashed lines the mean plus or minus one standard deviation. (B) Early in the experiment, people with greater apraxia showed particularly slow reach-to-use time with incongruent tools. A dot represents the performance of single LCVA individuals with their performance collapsed across all tools, with repetitions binned into pairs of trials across the task (red for incongruent tools and green for congruent tools).

We conducted a control analysis to check whether the effect of incongruence as a function of apraxia severity could be explained by the direction the hand needed to move to use the tool; that is, whether these differences could be explained by limb biomechanics (Flanagan and Lolley, 2001; Vindras et al., 2005; Cos et al., 2011). Specifically, congruent tools in our task required abductive, away-from-body movements while incongruent tools required adductive, toward-body movements. To address this concern, our task design included control trials in which the orientation of the tools was reversed, changing the required direction of use without affecting the incongruence of the tool (see Methods). Crucially, for reach-to-use time we did not observe a significant effect of hand-motion direction (χ^2^(1) = 1.07, p = 0.30) nor an interaction between hand-motion direction and Apraxia Severity (χ^2^(1) = 0.03, p = 0.87). This suggests that the direction of hand motion could not account for impaired planning with incongruent tools.

Finally, we assessed whether performance differed overall in the entire group of LCVA as compared to neurotypical controls. For RT, we observed a main effect of Incongruence (χ^2^(1) = 7.14, p = 0.007), again suggesting that incongruent tools required longer RTs than congruent tools (Incongruent tools: 552 ± 40 ms, Congruent tools: 498 ± 45 ms). We did not observe a main effect of Group (χ^2^(1) = 2.70, p = 0.10), nor an interaction between Incongruence and Group (χ^2^(1) > 0.27, p = 0.60). For reach-to-use time, we found a main effect of Group (χ^2^(1) = 7.73, p = 0.005), driven by prolonged reach-to-use time in people with LCVA compared to neurotypical controls (LCVA: 999 ± 50 ms, Neurotypical: 780 ± 38 ms). Neither the main effect of Incongruence (χ^2^(1) = 0.68, p = 0.41) nor the interaction between Group and Incongruence (χ^2^(1) = 2.67, p = 0.10) were significant. Thus, when considered as a whole, people with LCVA did not exhibit significant abnormalities in planning the use of incongruent versus congruent tools; rather, deficits were specific to individuals with a substantial degree of limb apraxia.

#### Impaired incongruent tool-use planning for people with apraxia is exacerbated early in exposure to novel tools

Anecdotally, we observed that some individuals with apraxia initially seemed to be uncertain about the direction they needed to push the tool handle to achieve the goal of moving the puck into the target. This observation raised the question of whether, across the six encounters with each tool, people with apraxia were able to improve their performance. To address this question, we explored the relationship between incongruence, apraxia severity, and learning in a post-hoc analysis. We examined reach-to-use time early, during, and late in the task by dividing trials into three bins and testing whether the interaction between incongruence and apraxia severity described above was also affected by repetition (i.e., bin). We found a main effect of Bin (χ^2^(2) = 66.55, p < 0.001), revealing that people with LCVA planned faster with more repetitions. A post-hoc pairwise analysis revealed that reach-to-use time was slower in the first bin (Bin 1: 1173 ± 65 ms) as compared to the two other bins (ps < 0.001, Bin 2: 944 ± 47 ms, Bin 3: 887 ± 50 ms). No difference was found between the second and third bins (p = 0.26). We also observed an interaction between Bin and Incongruence (χ^2^(2) = 7.23, p = 0.02), driven by longer reach-to-use time for incongruent tools than congruent ones specifically in Bin 1 (post-hoc test: p < 0.001, Bin 1: Incongruent Tool = 1253 ± 76 ms, Congruent Tool = 1081 ± 74 ms). No difference was found between congruent and incongruent tools for the other two bins (ps > 0.49). Crucially, we also observed an interaction between Incongruence, Apraxia Severity, and Bin (χ^2^(2) = 6.04, p = 0.04). This interaction indicates that reach-to-use time for incongruent relative to congruent tools was longer for people with more severe apraxia in the first bin (post-hoc test: p < 0.001, Fig. 2B). Interestingly, however, there was no significant difference as a function of Incongruence in the second or third bin (post-hoc tests: ps > 0.16). This suggests that people with more severe apraxia can improve in their ability to plan the use of incongruent tools, although it should be noted that their performance remained more impaired compared to less apraxic individuals within each bin (post-hoc tests: ps < 0.03). In contrast, people with less severe apraxia were not disproportionately slowed in planning the use of incongruent tools, and moreover did not improve their performance with experience (post-hoc tests: ps > 0.24).

#### Tool-hand incongruence slows the use of novel tools

Use of the novel tools was quantified by calculating average tool-use velocity. We observed a main effect of Apraxia Severity (χ^2^(1) = 5.76, p = 0.01), with tool-use velocity slowing as apraxia severity increased. The main effect of Incongruence was also significant (χ^2^(1) = 374.78, p < 0.001): tool-use velocity was slower for incongruent versus congruent tools (Incongruent tools: 69.40 ± 2.72 mm.s^-1^, Congruent tools: 125.60 ± 5.35 mm.s^-1^). A significant interaction between Apraxia Severity and Incongruence (χ^2^(1) = 13.78, p < 0.001) suggested that people moved faster when using congruent tools compared to incongruent tools particularly when they were less severely apraxic (post-hoc test: p < 0.001), although people with more severe apraxia remained significantly slower when using incongruent tools than congruent tools (post-hoc test: p < 0.001).

#### Tool-use planning deficits with incongruent tools may be associated with lesions in the praxis network

Our behavioral findings suggest that planning actions with incongruent tools is particularly challenging for people with limb apraxia following a left hemisphere stroke, especially when encountering a novel tool for the first time. In an exploratory analysis, we examined the locations in the brain that, when lesioned, were associated with poor planning of incongruent tool actions (i.e., longer reach-to-use time). Given the relatively small sample of participants (N = 29; Fig. 3A and 3B), this analysis is exploratory and should be interpreted with caution. Nevertheless, we observed that slower reach-to-use time for incongruent tools in Bin 1 was associated with lesions in the Superior Parietal Lobule (SPL), posterior Inferior Parietal Lobule (pIPL; including the posterior supramarginal gyrus and angular gyrus), and the posterior Temporal Cortex (pTC; including both posterior middle temporal gyrus and superior temporal gyrus) of the left hemisphere (p < 0.005 and cluster extent > 100 mm^3^; Fig. 3C and 3D; Table 2A); that is, posterior portions of the praxis network. When we ran the same analysis on the reach-to-use time for congruent tools, we found that impairments were associated with a substantially smaller cluster in the left lingual gyrus and an adjacent white matter tract, the left posterior thalamic radiation; p < 0.005 and cluster extent > 100 mm^3^; Fig. 3E and 3F; Table 2B). Finally, we ran a disjunction analysis to explore the brain areas associated with greater impairment for planning actions with incongruent as compared to congruent tools. We found that lesions primarily within the left pTC and pIPL were associated with greater planning impairments for incongruent compared to congruent tools (p < 0.005 and cluster extent > 100 mm^3^; Fig. 3G and 3H; Table 2C).

**Figure 3:**
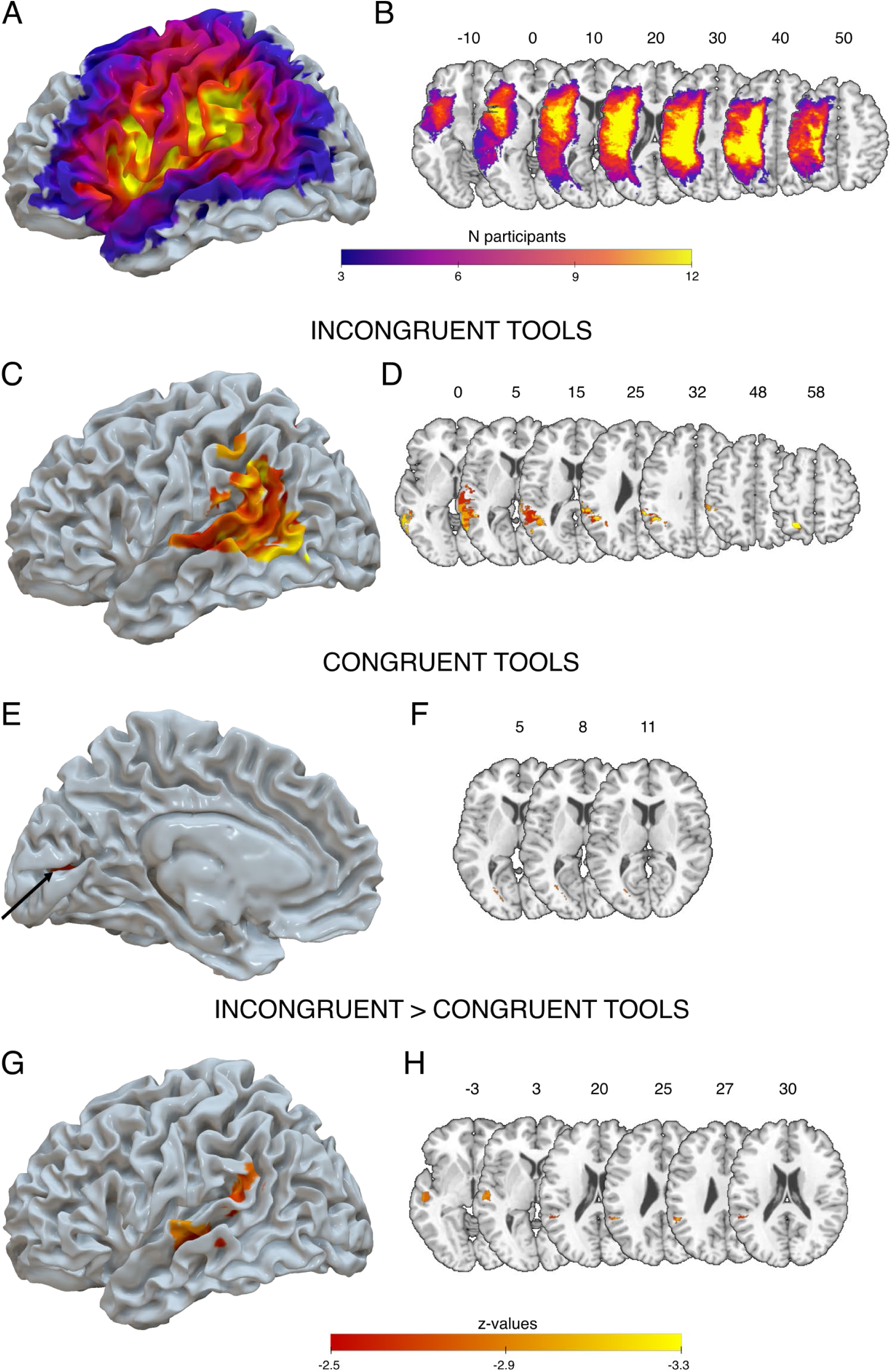
Tool-use planning deficits with incongruent tools may be associated with lesioned voxels in posterior portions of the praxis network. Inflated brain render (A) and axial-view brain slices (B) are shown for the lesion overlap map depicting voxels lesioned in at least 3 of the 29 participants; the color reflects the number of individuals for whom that voxel is lesioned (range: 3-12). Slice numbers reflect the corresponding z coordinates in MNI space. Thresholded maps shown on inflated brain renders of areas associated with slower incongruent tool reach-to-use time in the first bin with lateral view (C) and axial brain slices (D). Thresholded map of areas associated with slower congruent tool reach-to-use time in the first bin with medial view (arrow pointing to significant cluster) (E) and axial brain slices (F). Thresholded map of areas associated with greater incongruent than congruent tool reach-to-use times in the first bin with lateral view (G) and axial brain slices (H). A voxel was significant with a p-value < 0.005 one sided, corresponding to a z-value < -2.57 and if part of a cluster with at least 100 voxels (i.e., 100 mm^3^). All significant clusters are described in Table 1.

**Table 2:** Neural correlates associated with impaired tool-use performance: (A) Incongruent Tools, (B) Congruent Tools, (C) Incongruent Tools > Congruent Tools. Coordinates for the cluster centroid are in MNI space.

| *Brain area(s)* | *Cluster extent (mm<sup>3</sup>)* | *Peak Z-value* | *Cluster centroid coordinates* |
| --- | --- | --- | --- |
| **(A) Incongruent Tools** |  |  |  |
| Occipito-Parieto-Temporal Area | 10,360 | -3.72 | -55; -44; 14 |
| *Superior Temporal Gyrus* |  |  |  |
| *Middle Temporal Gyrus* |  |  |  |
| *Supramarginal Gyrus* |  |  |  |
| *Angular Gyrus* |  |  |  |
| *Inferior Occipital Gyrus* |  |  |  |
| *Middle Occipital Gyrus* |  |  |  |
| Superior Parietal Lobule | 343 | -3.54 | -22; -57; 57 |
| Supramarginal Gyrus | 222 | -3.72 | -56; -35; 47 |
| **(B) Congruent Tools** |  |  |  |
| Temporal Area | 174 | -2.88 | -24; -75; 9 |
| *Posterior Thalamic Radiations* |  |  |  |
| *Lingual Gyrus* |  |  |  |
| **(C) Incongruent Tools > Congruent Tools** |  |  |  |
| Temporal Area | 744 | -2.93 | -56; -21; 0 |
| *Posterior Superior Temporal Gyrus* |  |  |  |
| *Posterior Middle Temporal Gyrus* |  |  |  |
| Parieto-Temporal Area | 365 | -3.06 | -53; -44; 24 |
| *Superior Temporal Gyrus* |  |  |  |
| *Supramarginal Gyrus* |  |  |  |
| *Angular Gyrus* |  |  |  |

### Experiment 2: Effect of tool-hand incongruence on conventional tool-use gesture

In Experiment 1, we isolated the contribution of tool-hand incongruence to difficulties in action planning by examining how people use novel tools. Although we found a relationship between incongruence and apraxia severity, it is unclear whether tool-hand incongruence is relevant when people must use conventional and more familiar tools. The post-hoc analysis we conducted showed that even people with greater limb apraxia severity improved their planning of incongruent tools with experience, raising the possibility that the effect of incongruence would diminish as people become more familiar with a tool. Thus, in a second experiment involving the same group of people with LCVA from Experiment 1, we tested whether tool-hand incongruence affects the ability to gesture the use of conventional tools.

For each tool, we tested whether its degree of incongruence (as rated by a sample of neurotypical individuals) predicted Gesture-to-Sight accuracy (measured in the sample of people with LCVA regardless of their apraxia severity) along three movement components:

hand posture, arm posture, and amplitude/timing (see Methods). The analysis revealed a main effect of Tool-Hand Incongruence (χ^2^(1) = 19.42, p < 0.001), suggesting that across all individuals (regardless of apraxia severity), Gesture-to-Sight accuracy was worse for more incongruent tools. We also found a main effect of Gesture Component (χ^2^(2) = 152.13, p < 0.001), with people making more hand posture errors (Accuracy = 0.72 ± 0.03) compared to arm posture errors (Accuracy = 0.87 ± 0.02, post-hoc test: p < 0.001) or amplitude/timing errors (Accuracy = 0.89 ± 0.02, post-hoc test: p < 0.001), as we have shown previously (Buxbaum et al., 2003; Watson and Buxbaum, 2015). Importantly, we found an interaction between Tool-Hand Incongruence and Gesture Component (χ^2^(2) = 25.24, p < 0.001, Fig. 4). Post-hoc tests revealed that this interaction was driven by a relationship between arm posture and amplitude/timing errors with tool-hand incongruence (post-hoc tests: ps < 0.001), in which people committed more errors in these components as the tool-hand incongruence rating increased. No relationship was found between hand posture errors and incongruence ratings (post-hoc test: p = 0.15). These relationships remained even after controlling for ratings of tool familiarity. Taken together, these results suggest that tool-hand incongruence may indeed affect the ability of people with LCVA to gesture the use of conventional tools.

**Figure 4:**
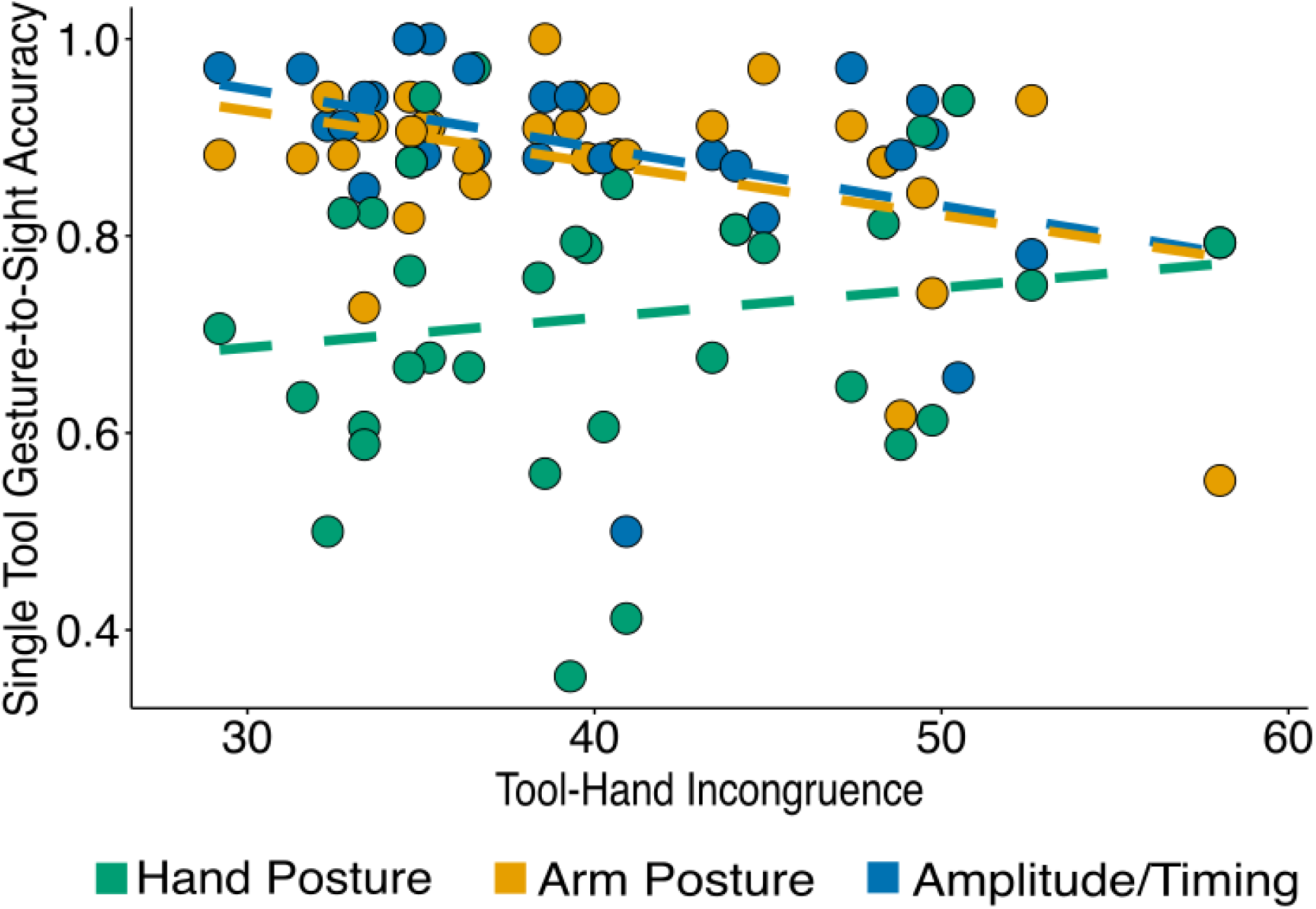
Item analysis of gesture-to-sight accuracy on individual tools as a function of tool-hand incongruence ratings for those tools. Accuracy of arm posture and amplitude/timing for participants was lowest for tools that were rated as incongruent, and highest for tools that were rated as congruent. Each dot represents a unique tool for which gesture-to-sight accuracy was averaged across individuals with LCVA and tool-hand incongruence was averaged across neurotypicals’ ratings. Gesture-to-sight accuracy was tested in people with LCVA and tool-hand incongruence was rated by an independent sample of neurotypical participants.

## Discussion

In a pair of experiments involving people with left-hemisphere stroke, we tested the hypothesis that a dissociation between tool and hand motion (i.e., incongruence) would give rise to action competition, increasing the difficulty of planning how to use both novel and conventional tools. In the first experiment, we demonstrated that individuals are more impaired at planning actions with novel incongruent compared to congruent tools, and that these impairments scale as a function of apraxia severity. This impairment was particularly pronounced early in the task, slowed people’s ability to execute actions with the tools, and was associated with lesions in the posterior inferior parietal lobule (pIPL) and posterior temporal cortex (pTC). In the second experiment, we showed that people with left-hemisphere stroke (regardless of apraxia severity) committed more errors when gesturing tool-use actions for conventional incongruent as compared to congruent tools. Errors arose primarily in components of the gesture (i.e., arm posture and amplitude/timing) expected to be sensitive to tool-hand incongruence, as opposed to hand-posture errors that are more associated with grasping the tool handle. Together, these findings support the hypothesis that incongruent tools invoke action competition between the motion of the tool tip and the opposing hand motion. After a LCVA, the ability to resolve this competition is presumably impaired, leading to longer planning times and impaired tool use.

How might competition between incongruent motions of the tool-tip and the hand arise? Many of the tools we use in daily life are congruent and can be viewed as an extension of the body (e.g., scissors, hammers, etc.). Thus, a movement congruent with the desired tool motion may be automatically activated when preparing a tool-use action (Beisert et al., 2010; Zhang et al., 2021). When the desired hand motion is incongruent with the tool motion, a second, incongruent motion may also become activated, resulting in two potential motions that compete for selection (Hardwick et al., 2019; Zhang et al., 2021). For these motions to compete, they must be sufficiently abstract (e.g., describing the desired motion path through space independent of whether the effector is a tool or hand, Kadmon Harpaz et al., 2014; Wong et al., 2016, 2019). Under typical circumstances, this competition is correctly resolved and the goal-directed response is performed. Our findings suggest that limb apraxia impairs the process of selecting between distinct tool and hand motions, similar to the observed deficits in resolving competition between representations associated with moving or using tools (Rounis and Humphreys, 2015; Watson and Buxbaum, 2015; Garcea et al., 2019). Interestingly, this disrupted competition resolution appears to be specific to tool use (e.g., unrelated to resolving competition in other cognitive tasks; Garcea and Buxbaum, 2023).

On the other hand, the neural correlates of tool-hand competition we observed (posterior SMG/IPL and pTC) are not identical to the more anterior regions previously associated with resolving competition between different representations associated with moving or using tools, i.e., the left anterior SMG, IFG, and insula (Watson and Buxbaum, 2015; Garcea et al., 2019; Garcea and Buxbaum, 2023). One possibility is that these distinct neural correlates within the ventro-dorsal stream (see Rizzolatti and Matelli, 2003; Binkofski and Buxbaum, 2013; Stoll et al., 2025) reflect different aspects of resolution of tool-action competition. That is, posterior regions (pTC and posterior SMG/IPL) within the ventro-dorsal stream may support an aspect of planning in which representations for tool versus hand motions are selected. This possibility is consistent with a number of studies indicating that the pTC is a hub for the representation of both hand and tool motions (Bracci et al., 2012, 2016; Perini et al., 2014; Lingnau and Downing, 2015; Pillet et al., 2024) and that the pIPL is involved in planning actions with incongruent tools (Gallivan et al., 2013; Ras et al., 2022). In contrast, more anterior regions (anterior SMG/IPL, IFG, and insula) may be involved in selecting the hand posture consistent with using versus moving the tool. The dissociation between these planning aspects is perhaps reflected in the findings from the second experiment, where we observed that errors when gesturing the use of incongruent compared to congruent tools occurred most frequently for the arm posture and amplitude/timing gesture components. In contrast, no such difference was observed for the hand posture component, which is known be sensitive to the competition between different “grasp-to-move” and “use” representations (Watson and Buxbaum, 2015). Action competition resolution may therefore help to select from among possible options to address different aspects of the task and determine which tool-related movement is ultimately produced.

Although our findings are consistent with impaired competition when selecting from among several potential actions, some previous studies examining tool-hand incongruence in neurotypical individuals have suggested that performance impairments might instead arise from difficulties in transforming the desired motion from tool-centered coordinates to hand-centered coordinates (Massen and Prinz, 2007; Beisert et al., 2010). This mapping from the desired target motion to hand movement (i.e., a reference-frame transformation) determines the proper hand motion associated with the tool and has been proposed to explain why incongruent tools are associated with longer reaction times compared to congruent tools (Massen and Prinz, 2007; Beisert et al., 2010). As other reference-frame transformations have been associated with the parietal lobe (e.g., Colby and Goldberg, 1999), it is possible that limb apraxia may disrupt the ability to compute this transformation, giving rise to slowed or impaired use of incongruent tools relative to congruent tools.

Note that these competition and reference-frame transformation hypotheses are not mutually exclusive, as both steps are critical for planning movements (Kim et al., 2021). However, as noted above, it is well-established across numerous studies that individuals with limb apraxia have difficulty resolving competition between “grasp-to-move” versus “use” actions and between different tool-use actions (Jax and Buxbaum, 2013; Rounis and Humphreys, 2015; Watson and Buxbaum, 2015; Garcea et al., 2019; Howard et al., 2019; Pizzamiglio et al., 2020; Rounis et al., 2021; Garcea and Buxbaum, 2023). In contrast, visuomotor transformation impairments have only very rarely been reported in these individuals (but see Mutha et al., 2010). Furthermore, the lesions typically associated with reference frame transformation impairments are bilateral and relatively more dorsal (dorso-dorsal) in the brain including the superior parietal lobule and dorsal premotor cortex (Perenin and Vighetto, 1988; Khan et al., 2005, 2013; Jax et al., 2009) compared to the lesions associated with apraxia or planning actions with tools, which are left-lateralized and more ventral (i.e., ventro-dorsal stream; (Binkofski and Buxbaum, 2013). Thus, when considered in the broader context of the existing literature (Watson and Buxbaum, 2015; Garcea et al., 2019; Garcea and Buxbaum, 2023), our findings are most consistent with an account emphasizing an impaired ability to resolve competition between potential tool-related actions. Future studies are needed to further test these hypotheses.

Despite initial impairments with using incongruent tools, we found that people with limb apraxia can learn to better plan their tool-use actions with increased exposure to tools. Indeed, people with greater apraxia severity exhibited longer reach-to-use times for incongruent tools early in the experiment, but by the end of the experiment the gap between incongruent and congruent tools vanished. These improvements might reflect an improved ability to resolve competition between discrepant tool and hand motions (i.e., improved action selection; e.g., Cisek, 2007; Botvinick et al., 2009; Jax and Buxbaum, 2010), perhaps via increased activation of the required (incongruent) hand motion with repeated exposure or by caching the required transformation between tool and hand reference frames (McDougle and Taylor, 2019). However, people with greater apraxia severity remained more impaired overall at planning the use of any tool (incongruent or congruent) compared to people with less apraxia severity, suggesting that learning cannot fully overcome the tool-use impairments associated with apraxia. Moreover, the second experiment suggests that incongruent tools remain challenging despite a great deal of prior exposure. Together, this suggests that for individuals with LCVA, short-term improvements with practice may reach a performance ceiling (Hardwick et al., 2017) and not be retained well – a hypothesis to be tested in future studies.

In summary, the data we have presented from two experiments indicates that tools associated with incongruent motions of the hand and tool-tip represent a particular challenge for action planning. This challenge likely arises from the need to resolve competition between representations of the potential motions of the hand and the tool. People with more severe limb apraxia seem particularly impaired at resolving this and other forms of tool-related action competition. Furthermore, competition resolution for different aspects of tool-use actions seem to rely on different portions of the ventro-dorsal stream. In particular, the posterior temporal cortex (pTC) and posterior inferior parietal lobule (pIPL) appear to be particularly important for resolving incongruence between hand and tool motions. Finally, despite initial impairments with incongruent tools, people with limb apraxia retain the capacity to reduce their tool use impairments with repeated practice. Overall, our study has shown that of the many factors that make tool use challenging (see Randerath et al., 2011), resolving the competition between tool-hand incongruence is critical for planning effective tool-use actions.

## Data Availability Statement

All relevant data, scripts and materials are available at https://osf.io/3965c/.

## Acknowledgments

We are grateful to Rand Williamson and Luke Carter for their support in recruitment, acquisition, and data preprocessing. We thank Apoorva Kelkar for preprocessing some of the brain lesions, H. Branch Coslett for checking the quality of both hand-drawn and automated lesion maps, John Krakauer for input in the conceptualization of the study, and Amanda Therrien for making available the Vicon setup for kinematics recording. This work was supported by NIH Grant #R01NS115862 to Aaron Wong.

