## Supplementary Materials for "Competition between tool and hand motion impairs movement planning in limb apraxia"

### Supplementary Text

*Constrained and unconstrained tool actions are equally difficult for people with apraxia to plan*

Our full set of tools included constrained tools (i.e., tools with a pivot) that enabled us to assess the effect of incongruence, along with unconstrained tools. Unconstrained tools may be more comparable to many of the tools used in daily life as they allow more degrees of freedom of manipulation, although because they lacked a pivot we were unable to design these tools to be incongruent. Nevertheless, we tested whether people planned constrained tool actions differently from unconstrained tool actions. We analyzed both RT and reach-to-use time for people with LCVA compared to neurotypical controls. For RT, we found a main effect of Constraint on RT ($\chi^2_{(1)}$ = 8.73, p = 0.003) revealing that constrained tools required longer RTs overall than unconstrained ones (constrained tools: 526 ± 42 ms, unconstrained tools: 478 ± 39 ms). However, we did not find a significant effect of Group ($\chi^2_{(1)}$ = 3.24, p = 0.07) nor an interaction between Group and Constraint ($\chi^2_{(1)}$ = 0.25, p = 0.61). For reach-to-use time, we found a main effect of Group ($\chi^2_{(1)}$ = 8.52, p = 0.003) revealing that people with LCVA were generally slower to reach to tools compared to neurotypical controls (LCVA: 993 ± 47 ms *vs.* Neurotypicals: 776 ± 32 ms). However, there was no main effect of Constraint ($\chi^2_{(1)}$ = 0.50, p = 0.47) nor an interaction between Constraint and Group ($\chi^2_{(1)}$ = 0.04, p = 0.83).

We also tested whether the severity of apraxia was related to any differences in using constrained versus unconstrained tools. For RT, we again found a main effect of Constraint ($\chi^2_{(1)}$ = 3.86, p = 0.04), with longer RTs associated with constrained tools (constrained tools: 571 ± 58 ms, unconstrained tools: 529 ± 54 ms). In contrast, we did not find an effect of Apraxia Severity ($\chi^2_{(1)}$ = 2.25, p = 0.13), nor an interaction ($\chi^2_{(1)}$ = 0.03, p = 0.85). For reach-to-use time, we found a main effect of Apraxia Severity ($\chi^2_{(1)}$ = 11.34, p < 0.001), but no main effect of Constraint ($\chi^2_{(1)}$ = 0.56, p = 0.45) nor an interaction between Apraxia Severity and Constraint ($\chi^2_{(1)}$ = 0.13, p = 0.71). Similar to the analysis with only constrained tools (see Results in Main Text), the main effect of apraxia severity reflected slower reach-to-use time for people with the most severe apraxia. Taken together, these results suggest that while people with LCVA are generally slower to plan tool-use actions, there were no consistent effects of constrained versus unconstrained (i.e., higher degrees-of-freedom) tools for people with LCVA relative to neurotypical controls.
